# Identification of a native Type I Secretion System cargo in *Zymomonas mobilis* and its application for extracellular enzyme secretion

**DOI:** 10.64898/2026.05.28.728472

**Authors:** Melissa Poma, Jose L. Munoz-Munoz, Ciarán L. Kelly

## Abstract

Engineering the ethanologenic Gram-negative bacterium *Z. mobilis* for the secretion of hydrolytic enzymes is a key step towards establishing a biofuel cell factory that uses complex waste material as feedstock. Secretion strategies in *Z. mobilis* have exclusively relied on signal peptides, which limit protein transport to the periplasm. To achieve single-step secretion across the *Z. mobilis* double-layered membrane, we sought to identify a native Type I Secretion System (T1SS) tag for fusion to proteins of interest. While a T1SS operon had been identified in the *Z. mobilis* genome, its native cargo had remained unknown and the use of T1SS secretion tags had so far been unexplored. Here, bioinformatic analysis identified the Major Intrinsic Protein (MIP) as a putative T1SS cargo, and its role validated through fusion of C-terminal sequences of two lengths (61 and 141 amino acids) to a heterologous β-galactosidase from *Bacteroides thetaiotaomicron*, expressed in *Z. mobilis*. The 141 amino acid tag, including two RTX domains, resulted in significantly higher secretion efficiency than the 61 amino acid tag lacking RTX repeats, consistent with the established role of RTX domains in preventing premature cytoplasmic folding, thus improving secretion. As extracellular secretion of hydrolytic enzymes has remained a major bottleneck in the development of *Z. mobilis* as a sustainable cell factory, the identification of a native T1SS secretion tag directly addresses this limitation, introducing a novel tool for enzyme delivery.

## Background

*Zymomonas mobilis* is a facultatively anaerobic, non-pathogenic, Gram-negative bacterium, characterised by an extraordinary ethanol production that reaches up to 98% of the theoretical maximum yield. Ethanol has emerged as a sustainable alternative to fossil fuels; however, its current bio-production strictly depends on the fermentation of simple sugars from crops, raising concerns about food security. As waste biopolymers such as cellulose, hemicellulose and lignocellulose are abundant, inexpensive and contain fermentable sugars, these feedstocks should drive future bio-ethanol manufacturing.

Attempts have been made to engineer *Z. mobilis* into a cell factory for the conversion of waste biopolymers into ethanol. Towards this goal, hydrolytic enzymes have been successfully expressed and produced, but their extracellular secretion has been poorly addressed, with efforts limited to the use of signal peptides ^1–5^, which target proteins to the periplasm rather than extracellularly. To ensure the transfer of proteins across the double-layered membrane, the use of bacterial secretion systems must be explored.

To date, at least eleven secretion systems have been identified in Gram-negative bacteria ^6^. These are commonly characterised by complex multimeric structures ^7^, challenging their application for molecular engineering purposes. Among the described systems, the Type I Secretion System (T1SS) stands out for its structural simplicity, comprising just three components: an ABC transporter, a membrane fusion protein (MFP), and an outer membrane protein (OMP) ^7^. While the genes encoding for the structural components and cargo of T1SS are often part of the same operon, the cargo or OMP can sometimes be found in separate *loci* ^8–10^. The translocation of T1SS cargoes occurs in an unfolded state, possibly to facilitate the transfer of different size passengers ^8^, which can span from 20 kDa to 1,500 kDa ^11^. These proteins are targeted to the T1SS translocator by a C-terminal secretion tag (29-60 aa) ^12^ that remains permanently attached to the cargo ^12^. T1SS secretion tags are reported to lack a consensus sequence ^13^, but conserved RTX (Repeats-in-ToXin) motifs can often be found upstream of the C-terminus ^12^. These are glycine- and aspartate-rich nonapeptides carrying the sequence GGXGXDXUX, where X can be any amino acid and U is a large hydrophobic residue ^12^. The RTX domains can reach up to 40 repeats ^14^, each separated by linkers of variable-length ^12^. Upon secretion, Ca^2+^ ions, which are predominant in the extracellular space, bind the RTX domains causing the protein to fold ^15,16^ (Fig. 1).

**Fig. 1.**
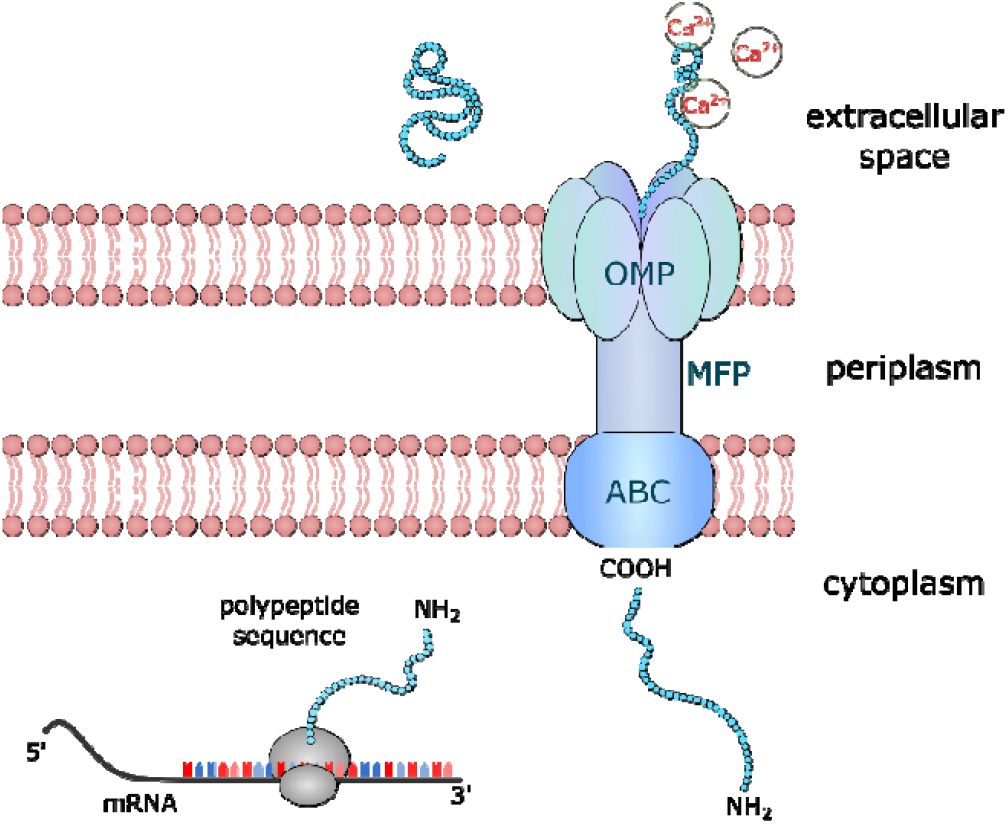
Type I Secretion System (T1SS). The T1SS translocon consists of an ABC transporter (ABC), a membrane fusion protein (MFP), and an outer membrane protein (OMP). The C-terminal region guides the secretion. Prior and during the secretion, the protein is maintained in an unfolded state; once the cargo reaches the extracellular space, Calcium ions (Ca^2+^), abundant extracellularly and nearly absent intracellularly, bind the protein triggering its folding.

The fusion of the C-termini of T1SS cargoes to heterologous proteins is a consolidated strategy to enable secretion in various Gram-negative bacteria ^13,17–19^. In the *Z. mobilis* genome, an operon including a T1SS OMP (*ZMO0253*), ABC transporter (*ZMO0254*), and MFP (*ZMO0255*) ^20^ has been identified, but its cargo is to date unknown. Identifying a native T1SS C-terminal could be key for the efficient secretion of proteins of interest in *Z. mobilis*.

In the present study, we sought to identify a native T1SS cargo in *Z. mobilis* as a source of a functional C-terminal secretion tag. Bioinformatic analysis of the genomic region flanking the *Z. mobilis* T1SS operon identified the Major Intrinsic Protein (MIP) as a candidate T1SS passenger, supported by the presence of RTX domains within its C-terminus and sequence similarity to known T1SS substrates. To validate this prediction experimentally, C-terminal sequences of two lengths were fused to a heterologous β-galactosidase from Bacteroides thetaiotaomicron and expressed in *Z. mobilis*, with extracellular enzyme activity used as a readout of secretion. This work establishes MIP as the first characterised native T1SS cargo in *Z. mobilis* and introduces its C-terminal sequence as a functional secretion tag for heterologous protein delivery.

## Results

### Computational Analysis Identifies MIP as a T1SS Cargo Protein in Z. mobilis

As cargoes of the T1SS are often located near the operon encoding the T1SS structural proteins, genes surrounding this genomic location were examined. There is a gene encoding for the cytoplasmic enzyme lactate-dehydrogenase located 115 bp downstream of the T1SS operon. Located immediately upstream of the *T1SS* operon, there is a gene annotated as encoding a Major Intrinsic Protein (MIP). The amino acid sequence of the encoded gene product was compared against known T1SS cargo proteins via the Protein BLAST^®^ database (non-redundant protein sequences database) ^23^, to look for similarity. The majority of the hits included Immunoglobin (Ig)-like domain-containing proteins, which are well-described in RTX adhesins, large multidomain T1SS passengers that participate in biofilm formation ^24^. Within the C terminus of the MIP, two RTX nonapeptides were identified, flanking another sequence that also resembles an RTX domain, although lacking two of the consensus amino acids (Fig. 2).

**Fig. 2.**
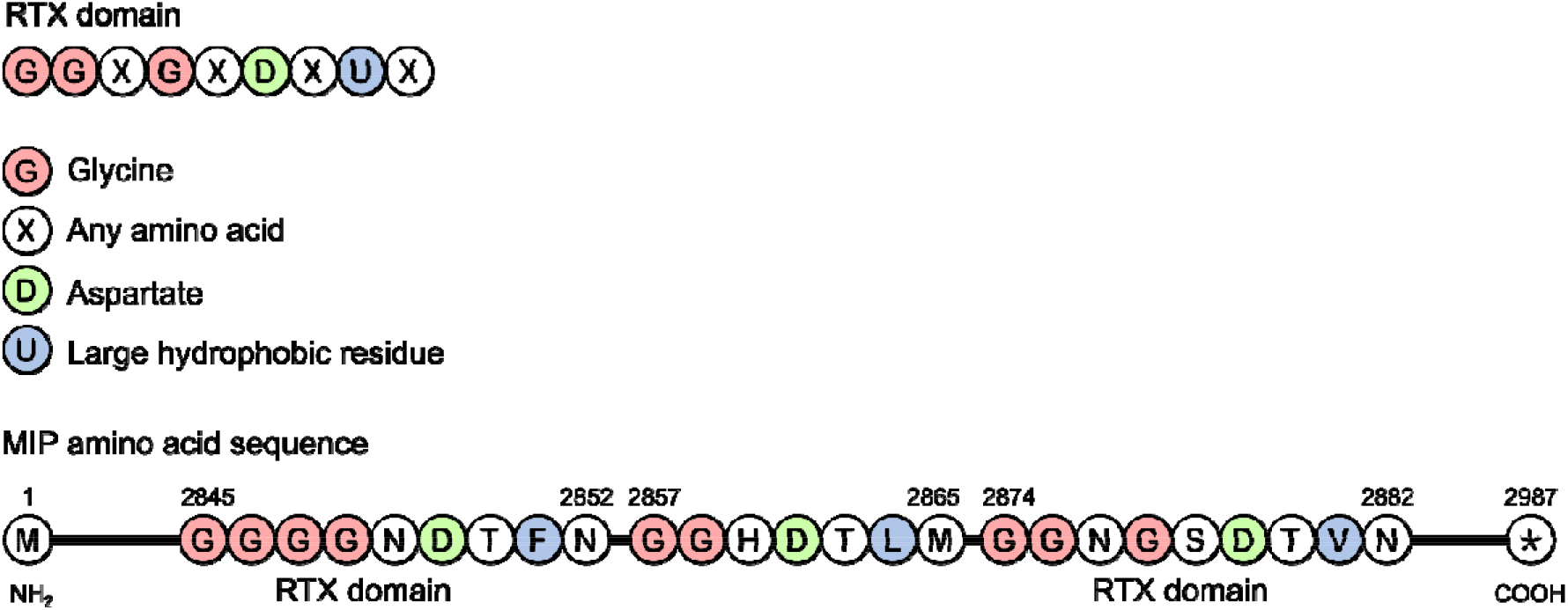
Repeats in toxins (RTX) domain composition and their arrangement within the MIP amino acid sequence. The RTX domain is a nonapeptide made of GGXGXDXUX, in which G is glycine, X is any amino acid, D is aspartate, and U is a large hydrophobic residue. In the MIP amino acid sequence of *Z. mobilis* there are two RTX domains. In the first RTX domain, (2845 aa – 2852 aa) and second RTX domain (2874 aa – 2882 aa), the large hydrophobic residues are respectively phenylalanine (F) and valine (V). In between these two conserved regions, there is a sequence that strongly resembles an RTX domain, except that it misses a glycine and any type of amino acid in fourth and fifth position.

This strongly suggested that MIP was a T1SS passenger protein, therefore the search for a T1SS secretion tag was narrowed to its C-terminus. Since known T1SS secretion tags are reported to range from 29 to 60 amino acids ^25–27^, a BLAST homology search using the last 60 amino acids of the MIP sequence was performed ^23^. 14 of the top 100 similarities were with T1SS C-terminal target domain-containing proteins, while 68 of them corresponded to Ig-like domain-containing proteins. The remaining hits included hypothetical proteins and domains found in proteins that are associated with biofilm production and reported to use T1SS for their secretion ^28–30^ (Fig. 3A) (Table S1). To further investigate this sequence for a potential secretion tag, the last C-terminal 60 amino acids of MIP were also run on the software HMMER, which performs fast and sensitive searches for protein homologies based on Hidden Markov Models (HMMs) ^31,32^. On HMMER, the five top hits for Bit score and E-value were annotated as T1SS C-terminal target domains. The remaining of the top 100 hits comprised C-terminal ends of proteins that are either unknown or described to be secreted through T1SS ^33–36^ (Fig. 3B) (Table S2). The five-best homologies with T1SS C-terminal tags were found in the last 53-55 amino acids of the MIP sequence, suggesting that the tag was likely within this region.

**Fig. 3.**
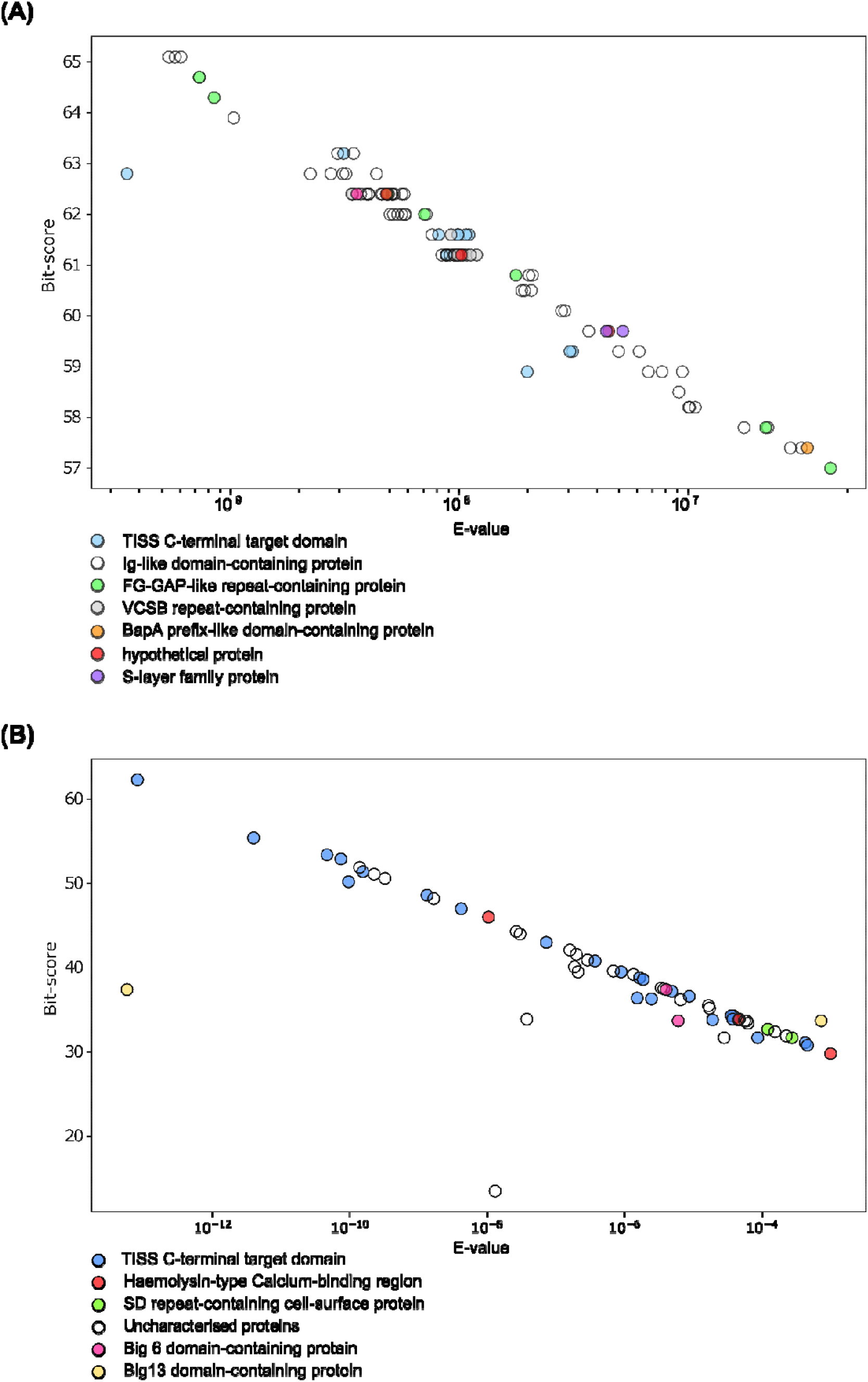
**(A)** Protein homologies of the 60 amino acids of the MIP C-terminus against the Protein BLAST^®^ database. The hits are plotted based on their Bit-score and E-value. The Bit-score represents the quality of an alignment, while the E-value indicates the probability that an alignment score was generated by chance. Most of the hits included Ig-like domain-containing proteins, of which MIP is predicted to be part. T1SS C-terminal target domains were the second most abundant hits. The remaining hits included: FG-GAP-like repeat-containing proteins, VCSB repeat-containing proteins, BapA prefix-like domain-containing proteins, hypothetical proteins and S-layer family proteins. FG-GAP repeats, VCSB repeats, BapA prefix-like domains, are found in proteins that participate in cell adhesion and biofilm formation ^37–39^. S-layer proteins are in the bacterial cell wall where they regulate the interactions with the environment and other cells ^308^. All these proteins are predicted to be secreted by T1SS ^29,30,37^. **(B)** Protein homologies of the last 60 amino acids of the MIP C-terminus identified by HMMER, which performs searches based on Hidden Markov Models. The hits are plotted based on their Bit-score and E-value. Most of the protein homologies were found with T1SS C-terminal target domains. The remaining hits included haemolysin-type Calcium-binding regions, SD-repeat-containing cell surface proteins, uncharacterised proteins, Big 6 domain-containing proteins and Big 13 domain-containing proteins. Haemolysin is a pathogenic protein secreted by T1SS. SD-repeats are found in surface proteins that participate in cell adhesion and biofilm formation ^34,35^. Big 6 and Big 13 are proteins containing Ig-like domains ^316^, of which the function and possible secretory pathway are unknown ^36^. The bioinformatic analysis on Protein BLAST^®^ and HMMER were done excluding the *Z. mobilis* genome from the search.

Besides MIP, we sought to identify more *Z. mobilis* T1SS cargo proteins as a source of alternative secretion tags. To this end, we scanned the *Z. mobilis* genome for RTX domains, due to their association with T1SS passenger proteins ^12^. The search was carried out by running the RTX domains of MIP on Protein BLAST^®^ against the *Z. mobilis* proteome database. This comparison identified a single sequence that could resemble an RTX domain, however, this motif was located in a periplasmic phosphate-binding protein (*ZMO1047*) and not repeated. Additionally, *ZMO1047* is predicted to have a N-terminal signal peptide ^40^, which is not found in T1SS cargoes, thus its participation in T1SS secretion must be excluded. As Ig-like repeats can also be indicative of T1SS cargoes, we searched for these domains across the whole *Z. mobilis* proteome using the software InterPro ^41^. A few proteins exhibited a single Ig-like domain, which however is not described to belong to T1SS passengers unless repeated.

### MIP C-Terminal Sequences drives β-Galactosidase Secretion in *Z. mobilis*

We tested whether the fusion of the C-terminal amino acids of MIP could result in the secretion of a heterologous β-galactosidase from *Bacteroides thetaiotaomicron* in *Z. mobilis*. The β-galactosidase gene was devoid of its native Sec signal peptide to avoid interference with the secretion aimed at T1SS, and the last 61 and 141 amino acids (aa) of MIP were fused to the C-terminus of the β-galactosidase. Unlike the 61 aa tag, the 141 aa tag included the RTX domains of MIP.

The two versions of T1SS-tagged β-galactosidases and a negative control lacking any tag were cloned downstream of the low-strength synthetic promoter P18 ^42^ in the broad-host range plasmid pMP002, generating the following plasmids: pMP109:β-galactosidase_Tag_61_, pMP110:β-galactosidase_Tag_141_ and pMP108:β-galactosidase. The plasmids were transferred into *Z. mobilis* by conjugation and the recombinant strains tested for secretion by measuring the extracellular β-galactosidase activity on the chromogenic substrate ortho-nitrophenyl-β-D-galactopyranoside (ONPG), which turns yellow (410 nm absorbance) upon cleavage. As *Z. mobilis* does not naturally possess a β-galactosidase or a lactose permease, which respectively degrade and up-take the ONPG analogue lactose, the degradation of the substrate can solely be attributed to the heterologous β-galactosidase in the extracellular space. Triplicate of recombinant strains were cultured statically at 30°C using 50 ml Falcon^®^ tubes containing 20 mL of ZMMG minimal medium. After 48h of growth, 100 µl of whole cultures were combined with 0.1 mM of ONPG and transferred to a 96-well microplate, which was incubated at 37 °C for an hour. When measuring absorbance at 410 nm in the plate reader, no change in colour was detected, indicating the lack of any active enzyme. As the *B. thetaiotaomicron* β-galactosidase is active at pH 7 ^43^, we measured the pH of *Z. mobilis* cultures and found it dropped from neutral to pH 3-4. This shift may result from static anaerobic growth, which produces lactic acid, acetic acid and CO_2_ ^97^. The experiment was repeated by replacing deionised water with pH 7 Phosphate buffer (PB) in the ZMMG medium. This stably maintained neutral pH throughout the bacterial growth, and a change in colour was finally detected when performing the enzymatic assay. *Z. mobilis* (pMP109:β-galactosidase_Tag_61_) and (pMP110:β-galactosidase_Tag_141_), showed a distinctively higher ONPG degradation than the negative control *Z. mobilis* (pMP108:β-galactosidase) (*p* value 0.018 and *p* value 5.7 × 10^-6^ respectively, two-sample *t*-test). Particularly, the presence of the RTX domains in pMP110:β-galactosidase_Tag_141_ led to a significantly higher ONPG degradation than its counterpart pMP109:β-galactosidase_Tag_61_ (*p* value 0.0007, two-sample *t*-test) (Fig. 4).

**Fig. 4.**
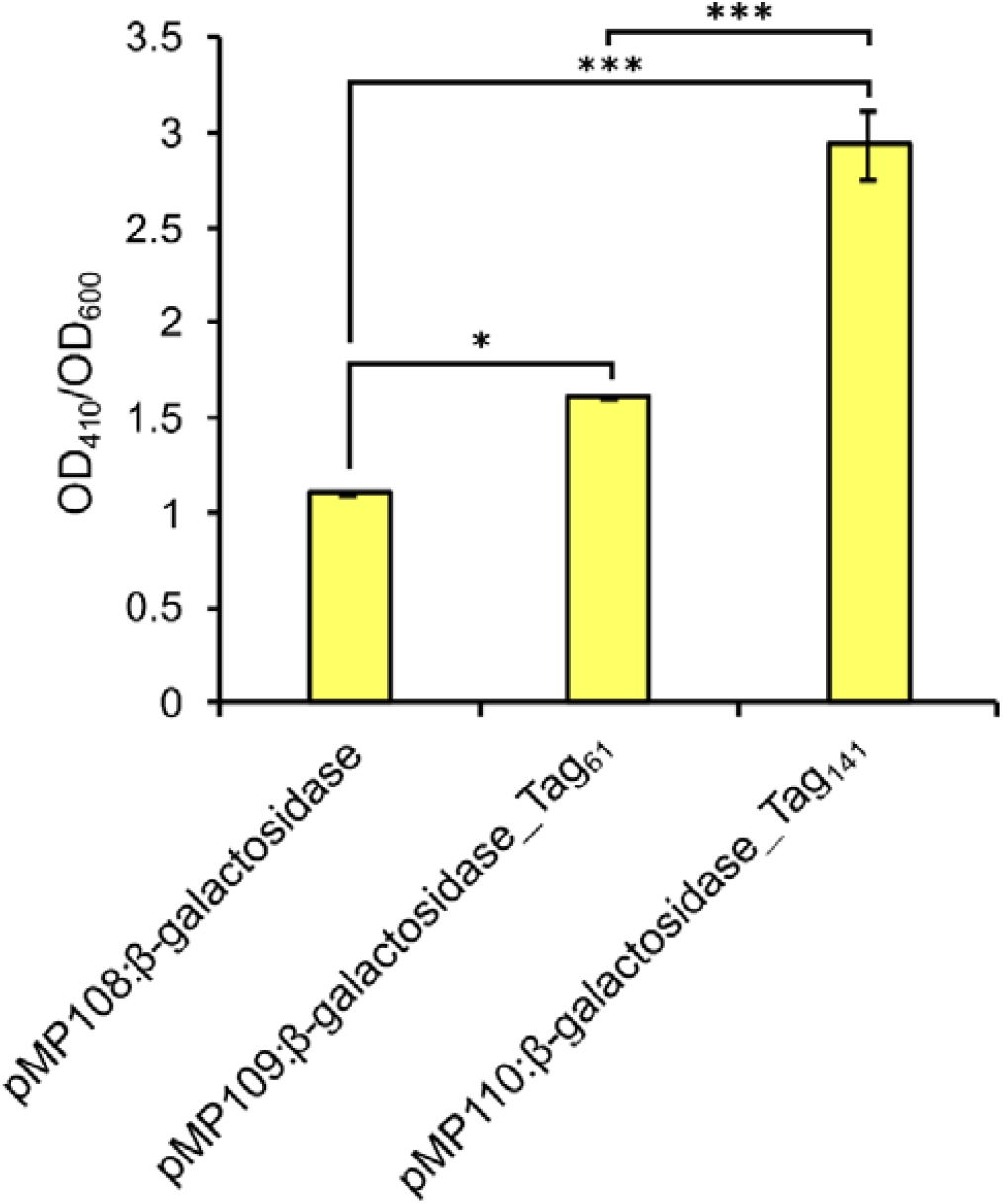
Secretion assay of a heterologous β-galactosidase from *Bacteroides thetaiotaomicron* in *Z. mobilis*. Secretion of the recombinant *Z. mobilis* strains (pMP108:β-galactosidase), (pMP109:βgalactosidase_Tag_61_), (pMP110:β-galactosidase_Tag_141_) was estimated by measuring absorbance at 410 nm, indicating the extracellular cleavage of the chromogenic substrate ortho-nitrophenyl-β-D-galactopyranoside (ONPG) by the heterologous β-galactosidase. The absorbance values at 410 nm were normalised by OD_600._ Samples were harvested during late exponential phase and absorbance at 410 nm and 600 nm measured using a microplate reader. *** = *p* < 0.01, * = *p* < 0.05, two-sample *t*-test. Error bars represent the standard deviation of three independent biological replicates.

## Conclusion

The use of secretion tags derived from T1SS passengers is a well-established strategy to achieve extracellular secretion of recombinant proteins in Gram-negative bacteria, bypassing the limitations of signal peptide-based approaches, which restrict protein transport to the periplasm. In *Z. mobilis*, native T1SS passengers and their application for recombinant protein secretion had not previously been investigated. Here, bioinformatic analysis of the *Z. mobilis* genome led to the identification of the Major Intrinsic Protein (MIP) as a T1SS cargo, supported by three independent pieces of evidence: its genetic proximity to the T1SS operon, the presence of RTX domains within its C-terminus, and sequence similarity to the C-termini of known T1SS substrates from other organisms. Notably, this similarity was detected within the last 60 amino acids of MIP, a region that does not include the RTX repeats, suggesting that conserved sequence features may exist among T1SS cargo C-termini beyond the RTX motifs. This is unexpected, as T1SS secretion tags are generally considered to lack a consensus sequence ^25,44,45^, and suggests that computational approaches may be more predictive than previously assumed.

Fusion of MIP C-terminal sequences of two lengths to a heterologous β-galactosidase revealed that the 141 aa tag, which includes the RTX domains, conferred significantly higher secretion efficiency than the 61 aa tag lacking them. This result is consistent with the established role of RTX domains in preventing premature cytoplasmic folding: protein folding rate is a critical determinant of T1SS secretion efficiency, with slowly folding or unfolded substrates showing enhanced secretion ^8,17,27^. The importance of maintaining cargo in an unfolded state prior to secretion is not unique to RTX-containing substrates, the non-RTX T1SS cargo HasA achieves the same outcome through binding to the SecB chaperone, which actively suppresses premature folding ^46–48^. This suggests that preventing premature folding is a common requirement across T1SS, regardless of the specific mechanism used to achieve it. The finding that even the shorter 61 aa tag, lacking RTX domains, still conferred higher secretion than the untagged control indicates that the functional C-terminal secretion tag resides within the last ~60 amino acids of MIP, with the RTX domains acting as a secretion-enhancer rather than an essential requirement for recognition by the T1SS translocon.

No additional T1SS cargo candidates were identified in the *Z. mobilis* genome through scanning for RTX domains or Ig-like repeats, suggesting that MIP may be the sole native T1SS passenger, but the existence of non-canonical cargoes that lack these features cannot be excluded. Most characterised T1SS typically secrete multiple substrates, for instance, the HasDEF and LipBCD systems of *Serratia marcescens* and the PrtDEF system of *Erwinia chrysanthemi* each translocate several passengers ^10^. A definitive catalogue of the *Z. mobilis* T1SS secretome would require proteomic comparison of wild-type and T1SS deletion strains. The polyploidy of *Z. mobilis* ^49^ currently represents a practical obstacle to generating complete knockouts across all genomic copies, and the inducible nature of T1SS cargo production in many organisms ^25^ suggests that identifying the conditions under which native passengers are expressed would require considerable additional investigation. In the absence of a complete secretome characterisation, heterologous T1SS tags from well-characterised systems in other Gram-negative bacteria represent a practical avenue for expanding the secretion toolkit in *Z. mobilis*, and screening these alongside the MIP tag across different cargo proteins would help determine how broadly these can be applicable in *Z. mobilis*.

The MIP secretion tag identified here provides a first route to extracellular enzyme delivery in *Z. mobilis*. Future work testing this tag with cellulolytic and other industrially relevant enzymes, including combinations of enzymes targeting complex biomass, will determine whether it can support the development of *Z. mobilis* as a consolidated bioprocessing strain capable of fermenting lignocellulosic feedstocks directly to ethanol.

## Experimental Procedures

### Bacterial strains and culture conditions

The bacteria used in this study are derivatives of the *Z. mobilis* ZM4 strain (NCBI accession number AE008692) with enhanced competence through deletion of *hsdSc hsdSp, E. coli* TOP10, and the diaminopimelic acid auxotrophic *E. coli* WM6026. The genetic details of these strains are reported in Table 1.

**Table 1.**
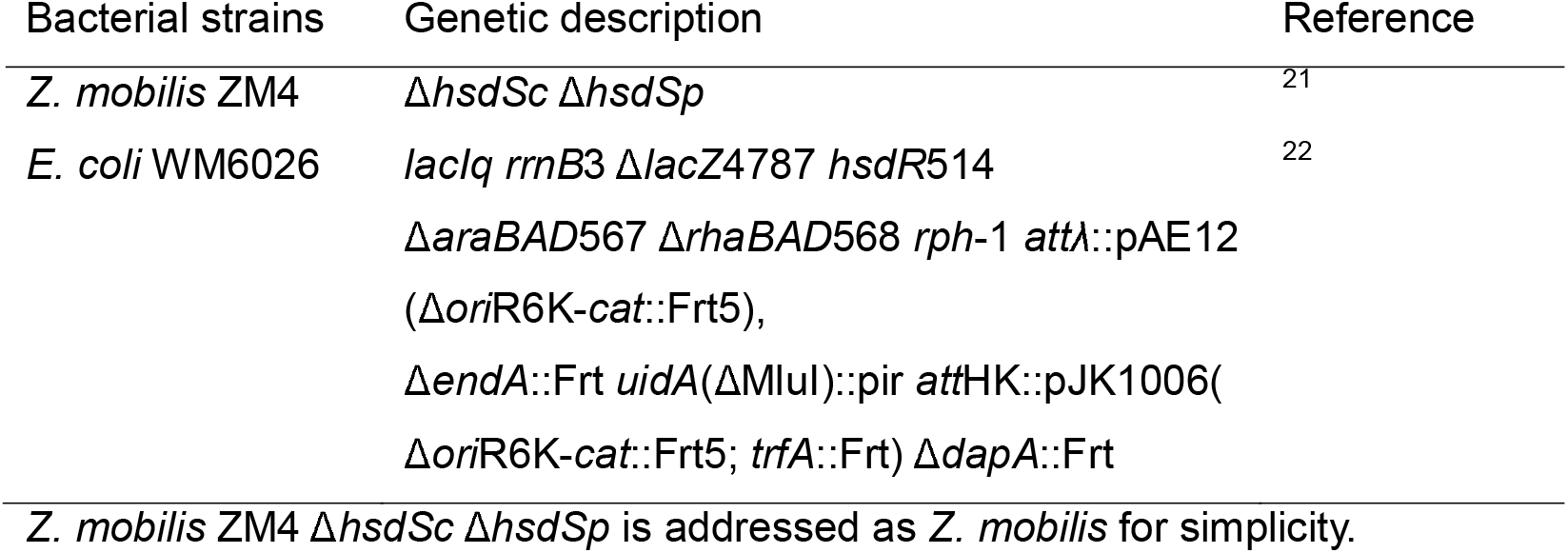
Bacterial strains used in this study.

| Bacterial strains | Genetic description | Reference |
| --- | --- | --- |
| *Z. mobilis* ZM4 | $\Delta hsdSc \Delta hsdSp$ | 21 |
| *E. coli* WM6026 | *lacIq rrnB3 $\Delta lacZ4787$ hsdR514 $\Delta araBAD567 \Delta rhaBAD568 rph-1 att\lambda::pAE12$ ($\Delta oriR6K-cat::Frt5$), $\Delta endA::Frt uidA(\Delta MluI)::pir attHK::pJK1006(\Delta oriR6K-cat::Frt5; trfA::Frt) \Delta dapA::Frt$* | 22 |

*Z. mobilis* ZM4 $\Delta hsdSc \Delta hsdSp$ is addressed as *Z. mobilis* for simplicity.

The medium used for growing *Z. mobilis* recombinant strains was *Zymomonas* Minimal Medium Glucose (ZMMG), containing a base solution of 1 g/L KH_2_PO_4_, 1 g/L K_2_HPO_4_, 500 mg/L NaCl, 1 g/L (NH)_4_SO_4_, and a trace elements solution of 200 mg/L MgSO_4_ x 7H_2_O, 0.010 g/L CaCl_2_ x 2H_2_O, 25 mg/L Na_2_MoO_4_ x 2H_2_O, 25 mg/L FeSO_4_ x 7H_2_O, 1 mg/L Calcium pantothenate (pH 6-6.5). To maintain neutral pH during growth, deionised water was replaced with pH 7 phosphate buffer in ZMMG. *E. coli* WM6026 22 was grown in LB supplemented with 0.1 mM diaminopimelic acid (DAP) at 37°C shaking at 200 rpm.

### Conjugation

Plasmids were transferred from *E. coli* WM6026 to *Z. mobilis* by conjugation. Mid-log cultures (OD_600_~0.5) of each strain were mixed 1:1, pelleted, resuspended, and spotted onto ZRMG agar supplemented with 10 g/L tryptone and 0.1 mM DAP. After overnight incubation at 30°C, cells were resuspended in ZRMG, incubated 2 h statically, and plated on ZRMG with the appropriate antibiotic.

### Construction of plasmids for testing the MIP C-terminus ability in secreting a heterologous β-galactosidase in Z. mobilis

The 60 and 141 amino acids of the MIP protein C-terminus (Table **2**) were fused to the C-terminus of a *Bacteroides thetaiotamicron* β-galactosidase (Accession number: UML59750) harboured in a pET28a vector. The gene had been previously deprived of its native Sec signal peptide, located in the first 19 amino acid of the β-galactosidase amino acid sequence.

**Table 2.**
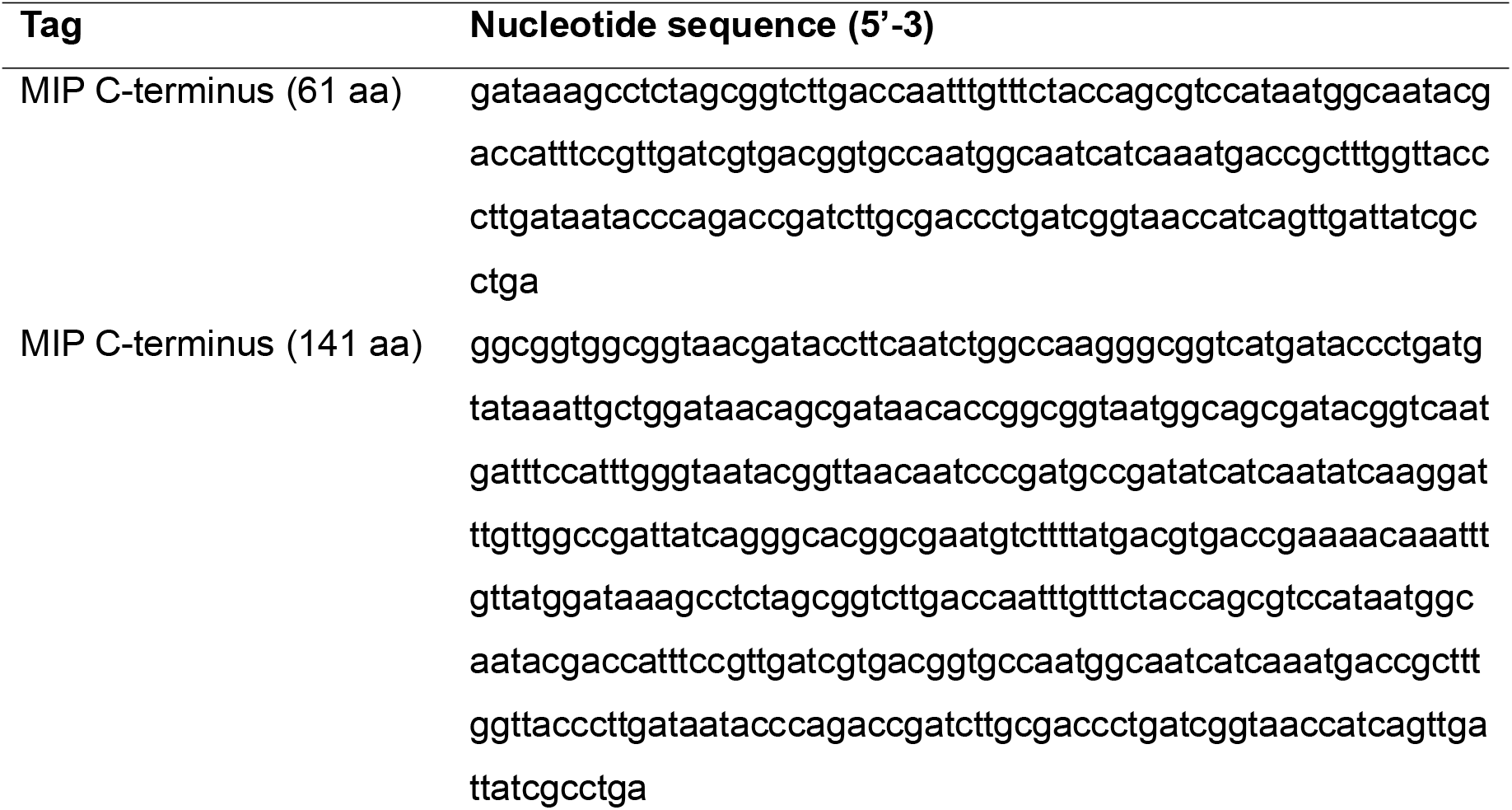
MIP C-terminus sequences used to tag the secretion of a heterologous β-galactosidase in *Z. mobilis*.

The regions including the pET28a RBS, β-galactosidase and T1SS tags were isolated and cloned in pMP037:P18:*sfGFP* in place of the *sfGFP* gene and its RBS. These assemblies generated the plasmids pMP109, pMP110 having the low-strength promoter P18 controlling the transcription of the β-galactosidase, respectively attached to the 61 and 141 amino acid MIP tags. As a negative control, the β-galactosidase without any secretion tag was cloned upstream of the P18 promoter, forming the plasmid pMP108. Plasmids are listed in Table 3.

**Table 3.** Plasmid list.

| Plasmid | Description |
| --- | --- |
| pMP002 | Broad-host-range plasmid containing pBBR-1 origin of replication, *sfGFP* gene, and spectinomycin resistance gene. |
| pMP108 | pMP002-derived plasmid, containing the low-strength promoter P18 upstream of the β-galactosidase gene from *B. thetaiotaomicron*, followed by the last 61 amino acid of the *Z. mobilis* MIP protein. |
| pMP110 | pMP002-derived plasmid, containing the low-strength promoter P18 upstream the β-galactosidase gene from *B. thetaiotaomicron*, followed by the last 141 amino acid of the *Z. mobilis* MIP protein. |
| pMP100 | pMP002-derived plasmid, lacking the *Z. mobilis* MIP C-terminus. |

## Resource availability

### Materials Availability

All plasmids generated in this study are available from the lead contact upon request.

### Data and Code Availability

All data supporting this study are available from the lead contact upon request.

## Author contributions

MP & CK designed study. MP led all experimental work. MP & CK wrote manuscript. JM helped with early study design and manuscript revisions.

## Acknowledgments

This work was funded by the EPSRC ReNU (Renewable Energy Northeast Universities) CDT at Northumbria University.

